# Follow that cell: leukocyte migration in L-plastin mutant zebrafish

**DOI:** 10.1101/2022.02.15.476948

**Authors:** J.B. Linehan, J.L. Zepeda, T.A. Mitchell, E.E. LeClair

## Abstract

Actin assemblies are important in motile cells such as leukocytes which form dynamic plasma membrane extensions or podia. L-plastin (LCP1) is a leukocyte-specific calcium-dependent actin-bundling protein that, in mammals, is known to affect immune cell migration. Previously, we generated CRISPR/Cas9 engineered zebrafish lacking L-plastin (*lcp1-/-*) and reported that they had reduced survival to adulthood, suggesting that lack of L-plastin might negatively affect the immune system. To test this hypothesis, we examined the distribution and migration of neutrophils and macrophages in the transparent tail of early zebrafish larvae using cell-specific markers and an established wound-migration assay. Knockout larvae were similar to their heterozygous siblings in having equal body sizes and comparable numbers of neutrophils in caudal hematopoietic tissue at two days post-fertilization, indicating no gross defect in neutrophil production or developmental migration. When stimulated by a tail wound, all genotypes of neutrophils were equally migratory in a two-hour window. However for macrophages we observed both migration defects and morphological differences. L-plastin knockout macrophages still homed to wounds but were slower, less directional and had a star-like morphology with many leading and trailing projections. In contrast, wild type macrophages were faster, more directional, and had a more streamlined, slug-like morphology. Overall, these findings show that in larval zebrafish L-plastin knockout primarily affects the macrophage response with possible consequences for organismal immunity. Consistent with our observations, we propose a model in which cytoplasmic L-plastin negatively regulates macrophage integrin adhesion by holding these transmembrane heterodimers in a ‘clasped’, inactive form and is a necessary part of establishing macrophage polarity during chemokine-induced motility.

## Introduction

Leukocyte plastin (L-plastin or LCP1) was first described as an abundant protein in chemically-transformed human fibroblasts (Goldstein et al. 1985; Lin et al. 1988). Further work established that this calcium-dependent actin-bundler was largely restricted to all normal leukocytes including neutrophils, monocytes, macrophages, T-cells, and B-cells (Lin et al. 1988; Lin et al. 1993; Pacaud and Derancourt 1993). As a result, L-plastin is now considered a key regulator of leukocyte chemotaxis, exocytosis, phagocytosis and other immune functions (Dubey et al. 2015); review by (Morley 2012). In addition to its role in the immune system L-plastin is found in many primary tumors and is correlated with clinical features of cancer, making the function of this protein highly relevant as a target for molecular therapies in immunology and oncology (Samstag and Klemke 2007; Shinomiya 2012; Ning et al. 2014; Tiedemann et al. 2019; Schaffner-Reckinger and Machado 2020).

Biochemically, L-plastin is thought to regulate the assembly of filamentous actin by stabilizing parallel bundles of F-actin strands (Ishida et al. 2017; Schwebach et al. 2017). These parallel bundles are a key component of extensible podia which are necessary for cell attachment and motility (Gardel et al. 2010; Leithner et al. 2016; Van Audenhove et al. 2016; Caswell and Zech 2018). Cell migration in solid tissues is also thought to depend on the cyclical binding of extracellular integrins (Chigaev and Sklar 2012). After integrin adhesion to the extracellular matrix (ECM), cells make actin-based protrusions at the leading edge, followed by de-adhesion and retraction of the trailing edge. In this context, L-plastin is a strong binding partner for the cytoplasmic tail of integrin β-chains in both resident immune cells and leukocyte cell lines (Arpin et al. 1994; Le Goff et al. 2010; Tseng et al. 2018). These findings implicate L-plastin as a novel component of ‘inside-out’ integrin signaling affecting cell adhesion and migration (Faull and Ginsberg 1996; Kolanus and Zeitlmann 1998(Faull and Ginsberg 1996; Kolanus and Zeitlmann 1998; Chigaev and Sklar 2012; Springer and Dustin 2012). However, much about L-plastin function is still unknown.

An important milestone in L-plastin research has been the construction of validated knockout mice (LCP1 -/-), allowing detailed study of immune function in these organisms. Although homozygous mutant mice are viable and fertile, they have numerous physiological defects in both innate and adaptive cell types including neutrophils, macrophages, T-cells, and B-cells (Wang et al. 2010; Todd et al. 2011; Morley 2013; Todd et al. 2013; Todd et al. 2016; Xu et al. 2016). However in mice, as in humans, not all aspects of immune development are easily accessible or visible *in vivo.*

L-plastin is highly conserved between humans, mice, and zebrafish. Unlike most zebrafish genes which are duplicated, there is only one L-plastin gene in this species, called *lcp1.* This gene produces a protein 84% identical and 93% similar to the human ortholog. The presence in zebrafish of a single, highly conserved L-plastin gene, plus a complete suite of all the myeloid lineages found in humans (Bennett et al. 2001; Crowhurst et al. 2002; Davidson and Zon 2004) provides an opportunity to dissect the role of this protein in specific cell types that are easily seen in the transparent embryos of this species.

Using CRISPR/Cas9 gene editing, we recently produced three independent zebrafish L-plastin knockout lines (Kell et al. 2018). Importantly, we have shown that homozygous mutant fish (*lcp1* -/-) express no detectable L-plastin protein at any point in the life cycle, whereas heterozygous and wild-type siblings express comparable endogenous amounts. All genotypes can mature and reproduce, allowing assessment of immune cell behavior throughout the life cycle. This paper describes the impact of L-plastin knockout on two types of innate immune cells – macrophages and neutrophils—in the early zebrafish embryo. We compare the cellular phenotypes observed to those of the existing knockout mouse model (LCP1 -/-), identifying areas of similarity, difference, and uncertainty. Finally, we propose a model in which L-plastin regulates integrin adhesion to regulate macrophage migration in solid tissues.

## Results

### Fish heterozygous and homozygous for knockout alleles of lcp1 have similar numbers of neutrophils at two days post-fertilization

L-plastin is one of the first markers of primitive hematopoietic cells in zebrafish and is expressed in myeloid precursors as early as 18 hours post-fertilization (Bennett et al. 2001; Ma et al. 2011). The role of L-plastin is to bundle F-actin, enhancing the stability of membrane protrusions during cell migration. Given this, we hypothesized that lack of L-plastin might cause reduced numbers or improper distribution of larval neutrophils, a key component of the early innate immune system. By two days post fertilization (2 dpf), neutrophils migrate posteriorly to colonize the zebrafish caudal hematopoietic tissue (CHT). The appearance of mature myeloid cells in this region marks a critical stage of hematopoiesis encompassing both cell multiplication and correct homing to key tissue niches in the organism. To assess the impact of L-plastin deficiency on these activities we analyzed CHT neutrophils in both heterozygous and homozygous mutant siblings (*lcp1* +/- vs. *lcp1* -/-) produced in a 1:1 ratio from a single pair of genotyped parents (**Fig. 1A**). On 2 dpf all embryos were fixed, stained for neutrophils using Sudan Black B, and then bisected into anterior and posterior segments. The anterior segment was processed for genotyping, while the posterior was mounted for imaging. From each image we measured both the total caudal fin area, representing body size (**Fig. 2A**) and the area stained with Sudan Black, representing neutrophils (**Fig. 2B**).

**Figure 1.**
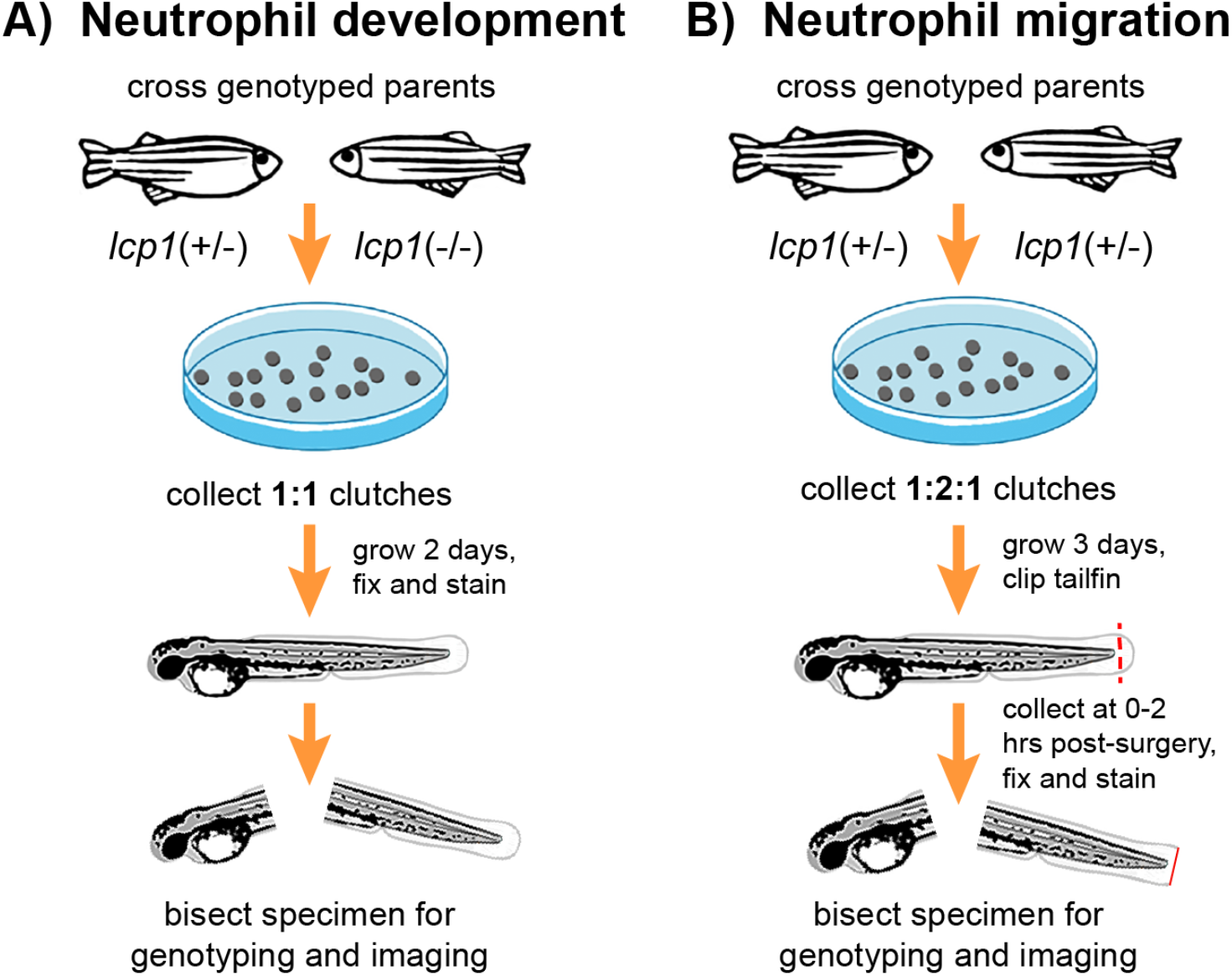
Flowchart of the experimental design. **A.** Neutrophil development experiments. **B.** Neutrophil migration experiments.

**Figure 2.**
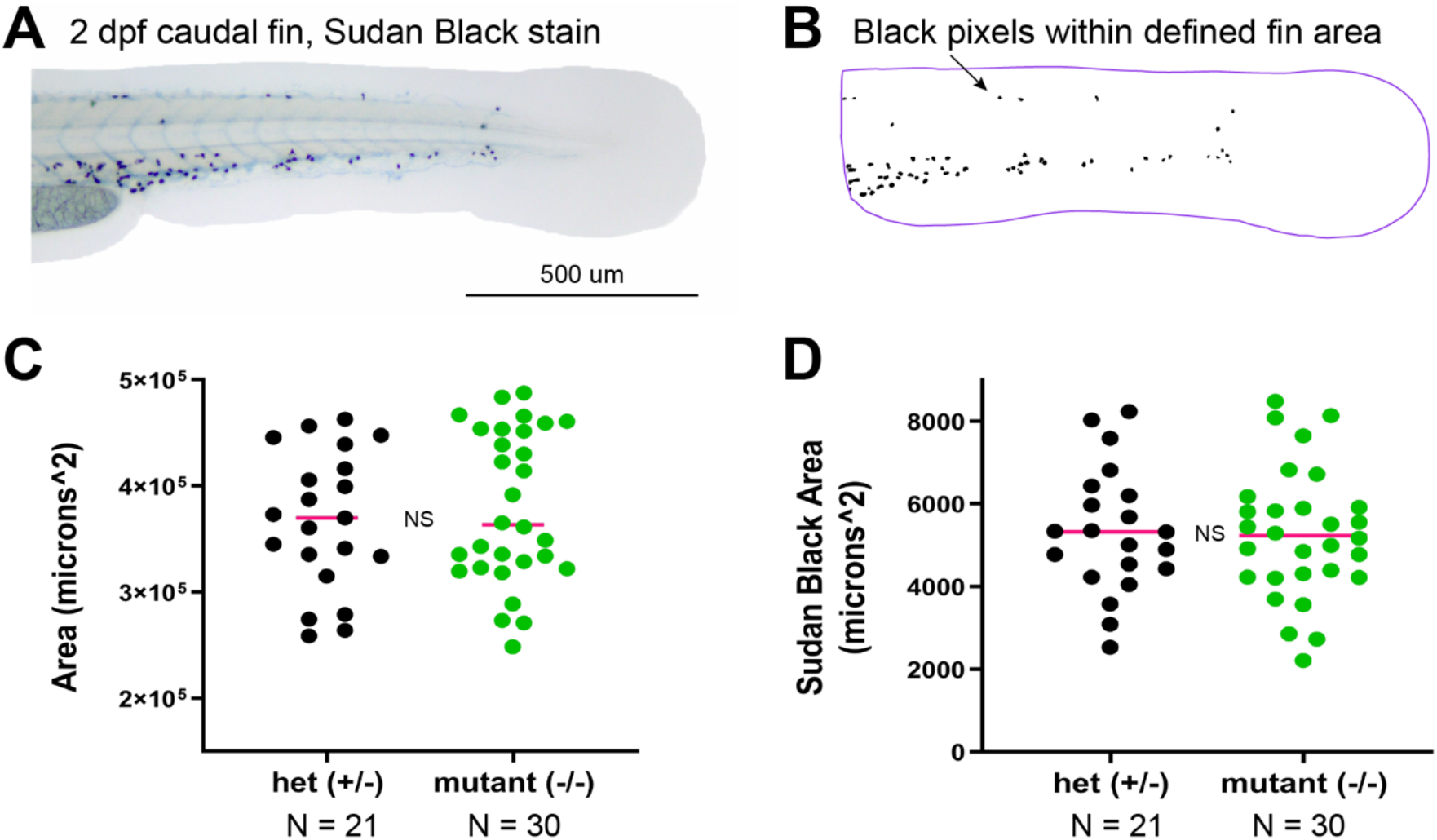
Neutrophil development in L-plastin knockout zebrafish. **A.** Flat-mount of a Sudan Black-stained larval caudal fin at two days post fertilization (2 dpf). **B.** Image analysis of the same specimen as in A. From a defined region of interest (purple outline) we extracted both the total fin area (= larval size) and the area of black pixels (= larval neutrophil population). **C.** Dot-plot of caudal fin areas for N = 21 *lcp1* heterozygotes (+/-, in black) and N = 30 *lcp1* homozygotes (-/-, in green). Magenta line = median value. **NS** = not significant. **D.** Dot-plot of Sudan Black-stained area for the same larvae analyzed in B. Magenta line = median value. **NS** = not significant.

After collecting several different clutches from the same mating pair, we processed 21 heterozygotes and 30 homozygous mutants in this assay. After calculating that the within-group variances were equal (F-test for variances, F = 1.209; df = [29, 20], p = 0.669) we then applied an unpaired, two-tailed t-test to the data. Using this test, we found no significant difference in caudal fin area between the two genotypes, indicating no size differential (**Fig. 2C**; t= 0.655, df = 49, p = 0.522). A similar test was used to compare the area of hematopoietic tissue stained with Sudan Black, representing neutrophils. On this variable we also found no significant intergroup variation (**Fig. 2D**; t = 0.128; df = 49; p = 0.898). In summary, L-plastin knockout zebrafish have normal-appearing neutrophil populations at 2 days post-fertilization. We infer that L-plastin protein is not essential for early neutrophil proliferation or localization to the caudal hematopoietic tissue.

### Lcp1-knockout does not impair wound-induced neutrophil migration

Continuing our study of neutrophils, we next stimulated wound-directed migration of these cells using an established larval finfold incision assay (Mathias et al. 2009; Roehl 2018). This experiment used multiple mating pairs in which both parents were heterozygous for the knockout allele, producing a 1:2:1 offspring ratio (**Fig. 1B**). After three days of development, batches of 15-20 sibling larvae were anesthetized and incised across the caudal finfold posterior to the notochord. All larvae in a batch were incised within 5 minutes. Incised larvae were euthanized and fixed either immediately after incision or after a 2-hour incubation at 28.5°C. After fixation, each whole larva was stained for neutrophils then bisected for processing, yielding a matched genotype and tail fin image of each specimen.

At zero hours post-surgery (0 hps) there were few if any wound-adjacent neutrophils. All genotypes tested had similar cell counts which never exceeded five cells per animal (**Fig. 3A and C**). In contrast, tail transection triggered neutrophil migration such that after two hours a typical specimen would have between two and ten cells within 100 microns of the amputation plane (**Fig. 3B and C**). However, the median number of migrating cells varied little among genotypes, being five, seven, and six neutrophils for wild type, heterozygous, and knockout larvae respectively. A nonparametric one-way ANOVA showed no significant effect of genotype on this value (Kruskal-Wallis test statistic H = 5.7, p = 0.06). We conclude that zebrafish neutrophils of any genotype (*lcp1* +/+, +/- or -/-) can make similar progress towards larval tail wounds within a 2-hour interval.

**Figure 3.**
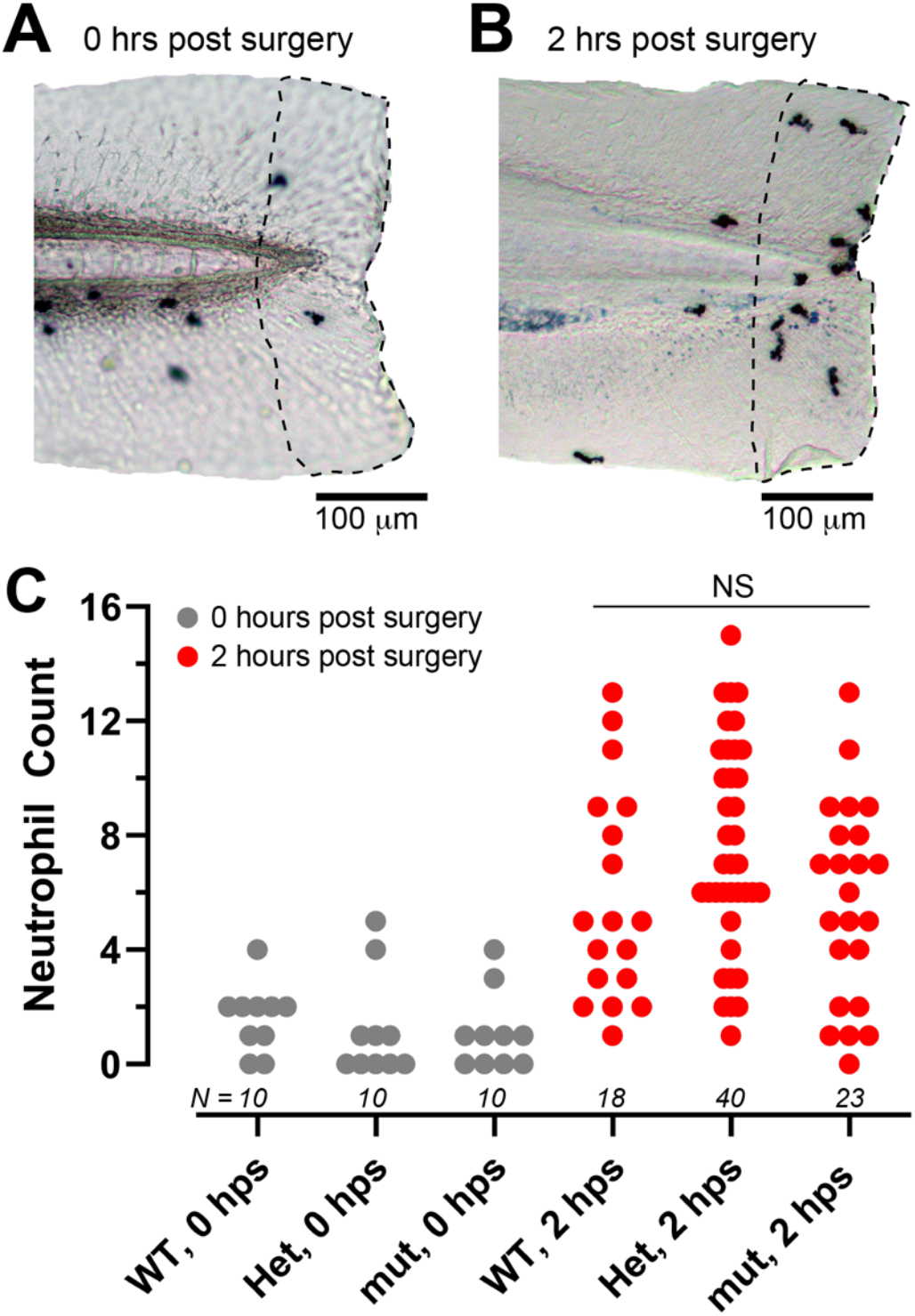
Neutrophil migration in L-plastin knockout zebrafish. **A.** Larval caudal fin fixed and stained immediately post-transection (0 hours post-surgery = 0 hps). Resident neutrophils were visualized with Sudan Black and manually counted in the region of interest (ROI) ~100 microns from the wound edge. **B.** Larval caudal fin fixed and stained 2 hours post-surgery (2 hps). Additional neutrophils are recruited to the region. **C.** Neutrophil counts for all 3 larval genotypes at 0 hps (gray) vs. 2 hps (red). **WT**= *lcp1* (+/+); **Het** = *lcp1* (+/-); **mut** = *lcp1* (-/-)

### L-plastin mutant macrophages are slower and less directional

Having assayed several aspects of larval neutrophils, we turned our attention to macrophages. To observe the migration of these cells in the context of wound healing, we repeated our larval finfold assay on two sets of embryos at 2 dpf: wild type embryos of a widely-used macrophage reporter line (*mpeg1.1*:EGFP) and L-plastin mutant embryos crossed into this line. One embryo at a time was wounded and fluorescent macrophages were filmed *in vivo* for two hours using the imaging parameters given in **Table 1**. After imaging, the target embryo was frozen for DNA analysis until the time lapse data was analyzed. To characterize macrophage behavior, the main variable measured was straight line displacement per minute (μ / min). Also called velocity, this is the speed of the cell as it moves between two successive image frames.

**Table 1.**
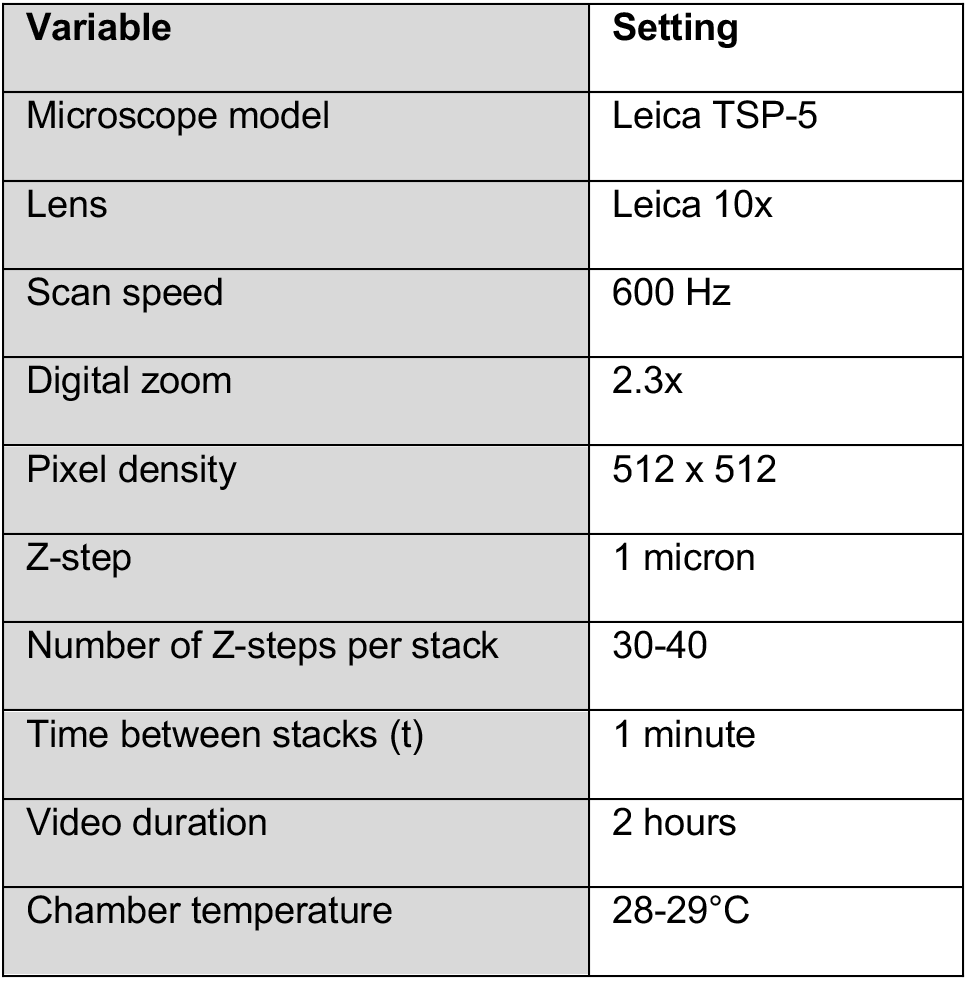
Image acquisition parameters for in vivo macrophage migration

| Variable | Setting |
| --- | --- |
| Microscope model | Leica TSP-5 |
| Lens | Leica 10x |
| Scan speed | 600 Hz |
| Digital zoom | 2.3x |
| Pixel density | 512 x 512 |
| Z-step | 1 micron |
| Number of Z-steps per stack | 30-40 |
| Time between stacks (t) | 1 minute |
| Video duration | 2 hours |
| Chamber temperature | 28-29°C |

Wound-directed macrophages were highly variable in speed, meaning that between successive frames they might move a small or large distance. However, in every wild type fish examined (N = 4) macrophages could consistently achieve speeds of more than 10 μ/min, with maximum speeds up to 30 μ/min (**Fig. 4A**). In contrast, all L-plastin fish examined (N = 3) had a truncated speed range. Almost all the steps were less than 10 μ/min, and none were above 20 μ/min, indicating that there were no extremely fast macrophages in mutant animals.

**Figure 4.**
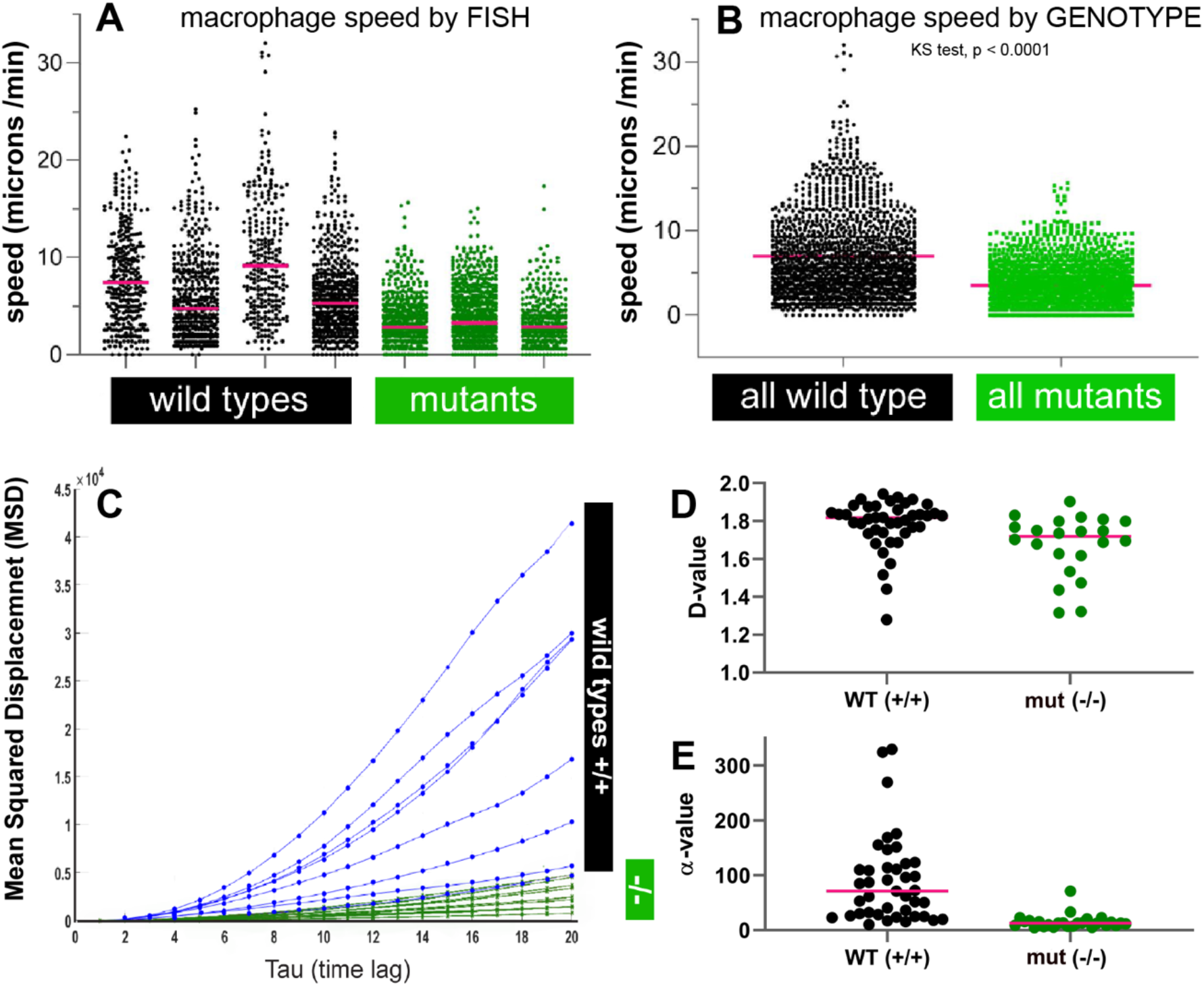
Macrophage migration. **A.** Dot-plot of macrophage speeds within individual wild type larvae (black) and homozygous mutant larvae (green) as measured during a two-hour *in vivo* fin-wounding assay. Each dot represents the distance traveled by a single cell between two successive image frames. **B.** Dot-plot of macrophage migration speeds for all wild type embryos combined (black)versus all mutant embryos combined (green). **KS** = Kolmogorov-Smirnov test for distribution form. **C.** Selected mean-squared displacement (MSD) curves for wild type and homozygous mutant cells over a 20-min tracking window (Tau range = 0 to 20). Wild type tracks (black) slope strongly upwards, indicating consistent directional progress during cell migration. In contrast, mutant tracks (green) have lower slopes, indicating less distance covered in the same period. **D** and **E.** Dot-plots of migration coefficients D and alpha, where D is a slope and alpha is an intercept. Wild type = black; homozygous mutant = green. Both groups have similar slopes (panel D) but very different intercepts (panel E). For definitions and calculations, see Methods.

To obtain a more precise estimate of macrophage speeds for each genotype we next pooled all wild type steps and all mutant steps, respectively. **Table 2** provides descriptive statistics for each pool. In this analysis, the median wild type (+/+) step was 8 μ/min, and the median mutant (-/-) step was 5 μ/s, a −35% speed reduction (**Fig. 4B**). Even more striking than the difference in median speed was the distribution of extremes, in particular the upper end representing the performance of the fastest-running macrophages. Using a sensitive non-parametric test for differences in the distribution form, we found that these two pooled datasets were not at all likely to differ in this key aspect due to sampling error alone (Kolmogorov-Smirnov test., p < 0.0001). We conclude that L-plastin deficiency reduces macrophage speed in response to larval tail wounds and nearly eliminates the cellular capacity for rapid bursts or ‘sprinting’.

**Table 2.**
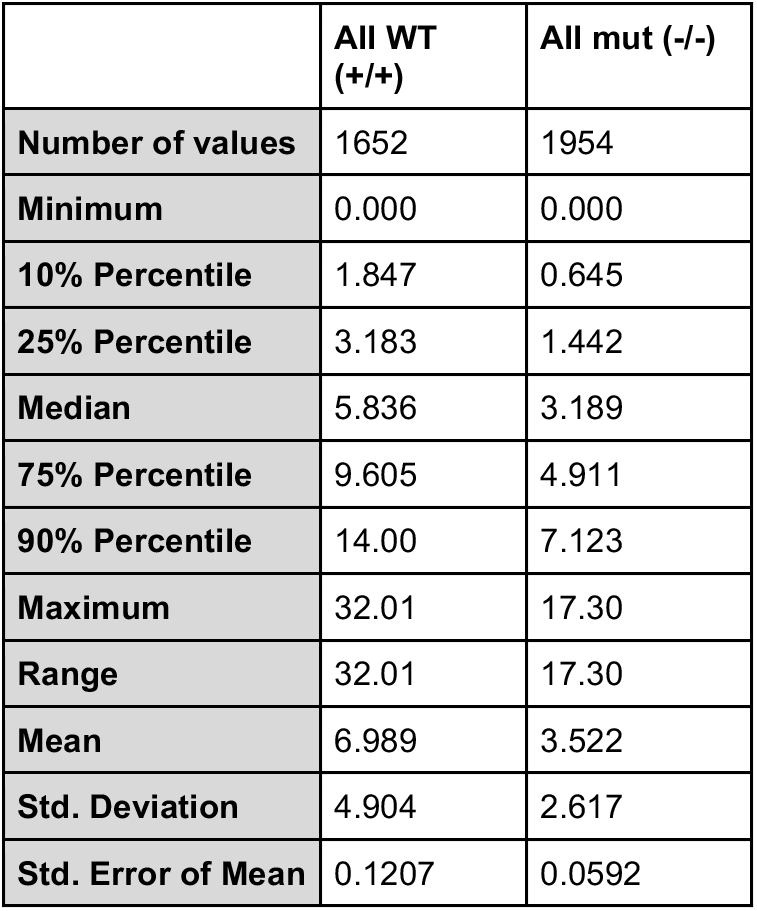
Pooled step analysis (microns/min) of wild type and lcp1 knockout macrophages

Whether a cell is slow or fast-moving, distance covered and directionality are critical to arriving at the site of injury or infection. The reduced ability of mutant macrophages to make single large steps means that they are limited to many small steps. Therefore they would be expected to cover less distance over the entire migration period (~ 2 hours). Confining our analysis to cell tracks that lasted more than 20 minutes (before becoming untraceable or reaching the wound edge), we calculated mean squared displacement (MSD) as an indicator of migratory persistence. In contrast to single-step displacement, which is instantaneous with respect to the time stamp, MSD provides a measure of the average cell displacement over increasing units of time. As such, MSD is sensitive to both the displacement of the macrophage and its directionality, providing additional data on how L-plastin deficiency affects cell locomotion. Although both wild type and mutant cells had upwards-sloping MSD curves, indicating positive chemotaxis, the MSD curves of wild type cells showed far greater upward curvature per time lag, indicating greater persistence.

Mathematically, a migrating cell’s MSD is useful because it can be compared to theoretical models of particle movement including diffusive random walks, physically-constrained random walks, or directed motion under an applied force (Codling et al. 2008; Krummel et al. 2016). For an applied force, the MSD function is modeled as the exponential function,

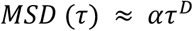

and a logarithmic transform of this equation yields the linear function log(MSD) = log(α) + D log (τ) where α is a scaling factor reflecting the material properties of the medium in which the cell moves. D is a constant quantifying the ‘super-diffusivity’ of the cell and depends on the probability of a displacement being a certain length. In this study α is considered a cell-specific parameter summarizing the macrophage response to wound stimuli. The diffusion coefficient D is considered to represent the total distance travelled by the cell, which is likely similar for all tracked trajectories. This is because *in vivo* all wound-responding macrophages are limited by the area of finfold tissue and in two hours some cells can reach the wound edge or ‘finish line’, thus limiting the total displacement they can achieve.

After extracting these parameters from tracks of the appropriate length we observed that both wild type (N = 41) and mutant tracks (N = 21) had similar distributions of D, meaning that both types of cells ‘diffused’ similarly in the 2-hour window. Both D-value distributions were predictably left-skewed, with the upper bound likely representing the limit of how far the cells could travel. In contrast, the a values reflecting the ‘barrier properties’ of the medium were very different between wild type and mutant cells. Mutant tracks yielded very low a coefficients, representing greatly reduced directional movement or ‘drive’. Wild type tracks yielded a coefficients that were highly variable but regularly exceeded 100-200x of mutant tracks, reflecting highly effective penetration through the ECM whilst traveling towards the goal.

Consistent with these quantitative analyses, the slower pace of mutant macrophages was reflected in how many reached the wound edge in the space of two hours and how they behaved once reaching this goal. Although it was not possible to accurately count or track individual macrophages close to the wound edge (due to the lower resolution of the images collected *in vivo* and the superimposition of cell boundaries after Z-projection) the following general trends were observed. In wild type larvae the fastest macrophages reached the wound edge very early, clustering there as more cells arrived (**Fig, 5**). These earliest-arriving cells were then observed to meander, reverse, and initiate retrograde migration. In mutant animals the fastest macrophages did home to the wound and meander, but fewer cells accumulated and there was no observable reverse migration in the two-hour time window.

**Figure 5.**
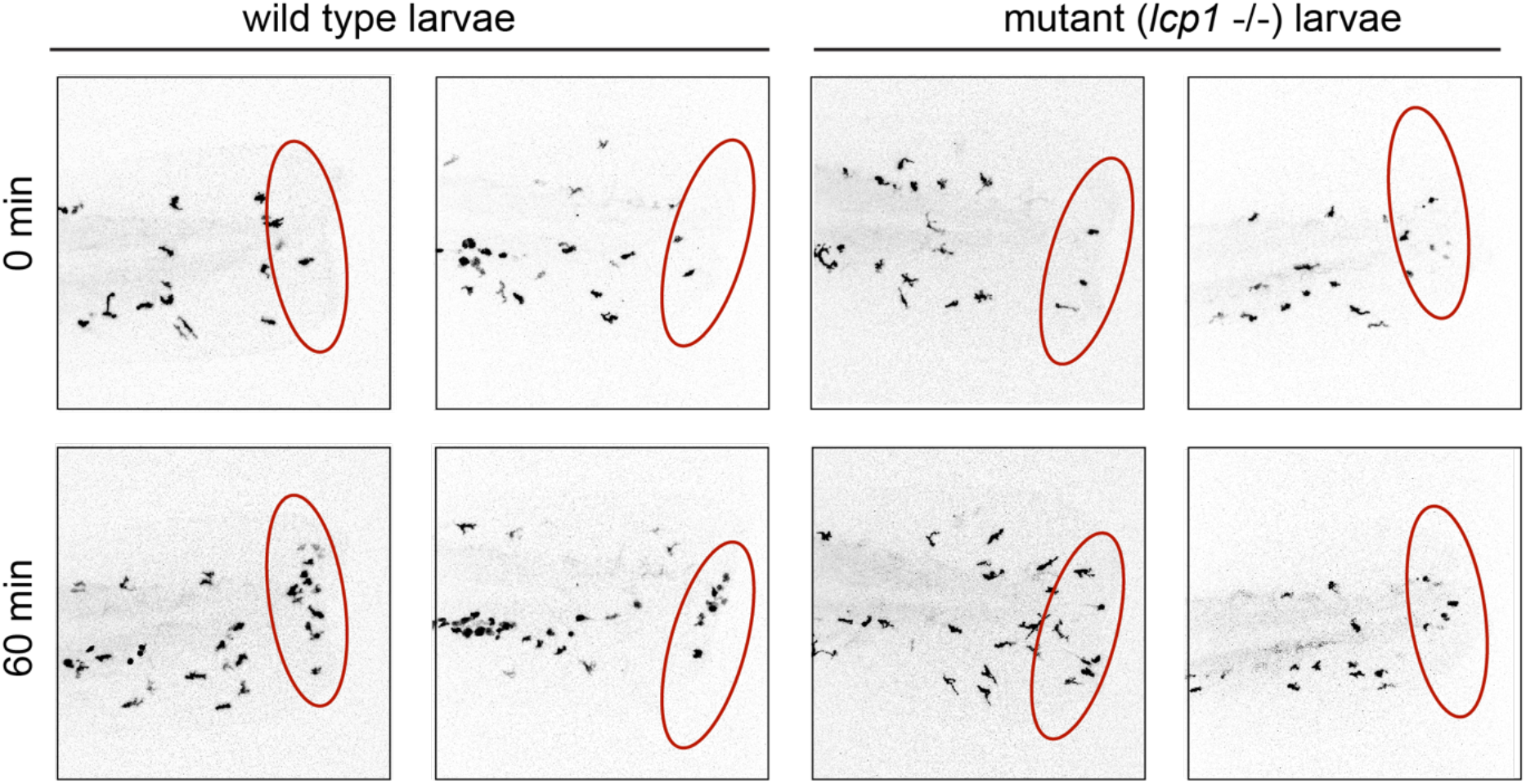
*In vivo* macrophage migration in wild type and L-plastin knockout larvae. Top row = cell distribution at the start of image capture (0 min). Bottom row = cell distribution 60 min later. Wild type larvae. (first two columns) accumulate more cells at the wound margin, approximated by the red ellipse.

Finally, in addition to cell speed, directional persistence, and reverse migration we observed phenotypic differences in cell morphology. Notably, most migrating mutant macrophages appeared highly ramified or ‘star-like’, with many leading and trailing projections (see **Supplemental File - Macrophage Tracking Movie**). In contrast, most migrating wild type macrophages had few or no projections, adopting a more streamlined, ‘slug-like’ morphology.

## Discussion

Many proteins bundle actin (Amos and Amos 1991). However, these tend to be conserved throughout eukaryotes and expressed in all cell types. L-plastin is unique because it is restricted to vertebrates and is normally expressed only in immune cells (Lin et al. 1993; Arpin et al. 1994). This suggests that L-plastin evolved from ancestral members of the plastin/fimbrin family but became specialized in vertebrates for immune cell function.

The first vertebrates to evolve an immune-cell repertoire were fishes, of which the zebrafish is a well-studied example. Over the last two decades zebrafish have become a central experimental model for studying immune cell development, migration, and behavior (Traver et al. 2003; Trede et al. 2004; Renshaw and Trede 2012). However, most information on L-plastin comes from humans and mice, whereas the corresponding protein in zebrafish has received less attention. Here we summarize our findings from L-plastin knockout fish (*lcp1* -/-) and compare these with cell-specific phenotypes in the existing mouse model (Chen et al. 2003).

Our principal finding is that *lcp1* knockout macrophages appear both migrationally deficient and morphologically distinct in the context of wound-directed migration. Mutant macrophages can still home to wounds but are incapable of reaching peak speeds on their journey and are less persistent as measured by mean-squared displacement (MSD). Although this effect was quite strong our experiment is limited in two ways. Only a small number of larvae were tracked *in vivo* (N = 7) and the wild types and transgenics were not full siblings but derived from different lines. Therefore we cannot rule out a line-specific effect or similar confounding factor influencing the differences observed. Such results, although intriguing, should be considered tentative until similar work is performed using larger numbers of larvae and full-sibling controls.

The impaired macrophage migration seen in our zebrafish tail-wounding experiments parallels in part the biology of experimental infections in L-plastin knockout mice. Specifically, when challenged with inhaled *Streptococcus pneumoniae,* up to 80% of LCP1 -/- mouse pups die, compared to ~8% of their wild type littermates (Deady et al. 2014). Using flow cytometry on lung aspirates, it was determined that knockout pups had fewer macrophages in their lung alveoli, and that this was a consequence of defective macrophage migration and not phagocytosis ability (Todd et al. 2016). It was concluded that L-plastin deficient macrophages cannot reach the lung in large numbers, leading to reduced bacterial clearance and rapid death. In this report we show that in zebrafish L-plastin deficient macrophages are less able to migrate to tail wounds. However, we did not study phagocytosis capacity or resistance to infection. Also unknown is the immune status of our L-plastin knockout fish later in the life cycle. Although zebrafish macrophages are far less visible at adult stages, these cells can be observed *in vivo* in thin tissues such as fins (Petrie et al. 2014), or *ex vivo* after blood or body cavity collection (Afonso et al. 1998). In future studies such methods might be used to assess macrophage phenotypes in adults using our previously-generated L-plastin mutant fish lines.

A qualitative observation from our study is that, in zebrafish, L-plastin mutant macrophages appear to migrate as ‘dendritic’ cells with many persistent, long projections. To the best of our knowledge this is the first time that the shape of migrating macrophages has been measured in a L -plastin deficient vertebrate *in vivo.* What causes this morphology is not known. However, based on leukocyte studies in mammals, we speculate that it is related to podosome instability and/or misregulation of integrin adhesion. A podosome, or ‘foot organ’, is an adhesive protrusion used by macrophages and other migrating cell types (Buccione et al. 2004; van den Dries et al. 2019). Recently, macrophages from wild type mice and LCP1 -/- mice were examined *in vitro* using immunohistochemistry and time-lapse microscopy (Zhou et al. 2016). In wild type macrophages F-actin and LCP1 were co-localized in podosomes. LCP1 -/- mouse macrophages still formed podosomes, but the stability of F-actin in these structures was decreased and individual podosomes were less persistent. Although not tested in zebrafish, this mechanism could explain the link between L-plastin deficiency and altered cell morphology.

In addition to its role in stabilizing actin-rich cell protrusions, L-plastin is a cytoplasmic binder of integrin tail domains. In this role, it extends its reach is beyond actin to regulate cell adhesion to the ECM. The outside part of integrins can have several configurations of which ‘clasped’ has the lowest affinity and ‘extended’ the highest (Askari et al. 2009). Overall, the balance of these configurations influences both cell adhesion and cell signaling (Kolanus and Zeitlmann 1998; Luo et al. 2007; Bachmann et al. 2019). In 2010, L-plastin was identified as a binding partner of the integrin β-chain and colocalized with integrins in immune cells (Le Goff et al. 2010). In 2018, an unbiased pull-down screen for cytosolic partners of clasped leukocyte integrin αMβ2 identified human LCP1 as one of the strongest-binding partners (Fig. 6 Tseng et al. 2018). Using human leukocyte cell lines, this latter study then knocked down LCP1 and found that L-plastin depleted leukocytes were more adhesive, compared to controls, under conditions of flow. Taken together, these independent findings place L-plastin as a novel cytoplasmic suppressor of hematopoietic integrin adhesion operating through and ‘inside-out’ signaling mechanism (Nolte and Margadant 2020).

Extending these findings to our zebrafish observations, **Figure 6** illustrates a possible mechanism of action. In wild type macrophages L-plastin negatively regulates macrophage adhesion by holding transmembrane integrin heterodimers in a ‘clasped’, inactive form (**Fig. 6A**). L-plastin knockout reverses this effect, converting clasped integrins to an unclasped, active form (**Fig. 6B**). This shift in the balance between active and inactive integrins makes mutant cells constitutively ‘sticky’, slowing their progress and changing their morphology. Although able to extend many projections, they may be less able to detach them, resulting in dendritic morphology and slowed migration. Additional observations are needed to confirm this model and it will be complex to navigate the tremendous variance of current experimental platforms, which differ in the choice of species, organ system, developmental stage, cell type, and assay (*in vivo* vs. *in vitro*).

**Figure 6.**
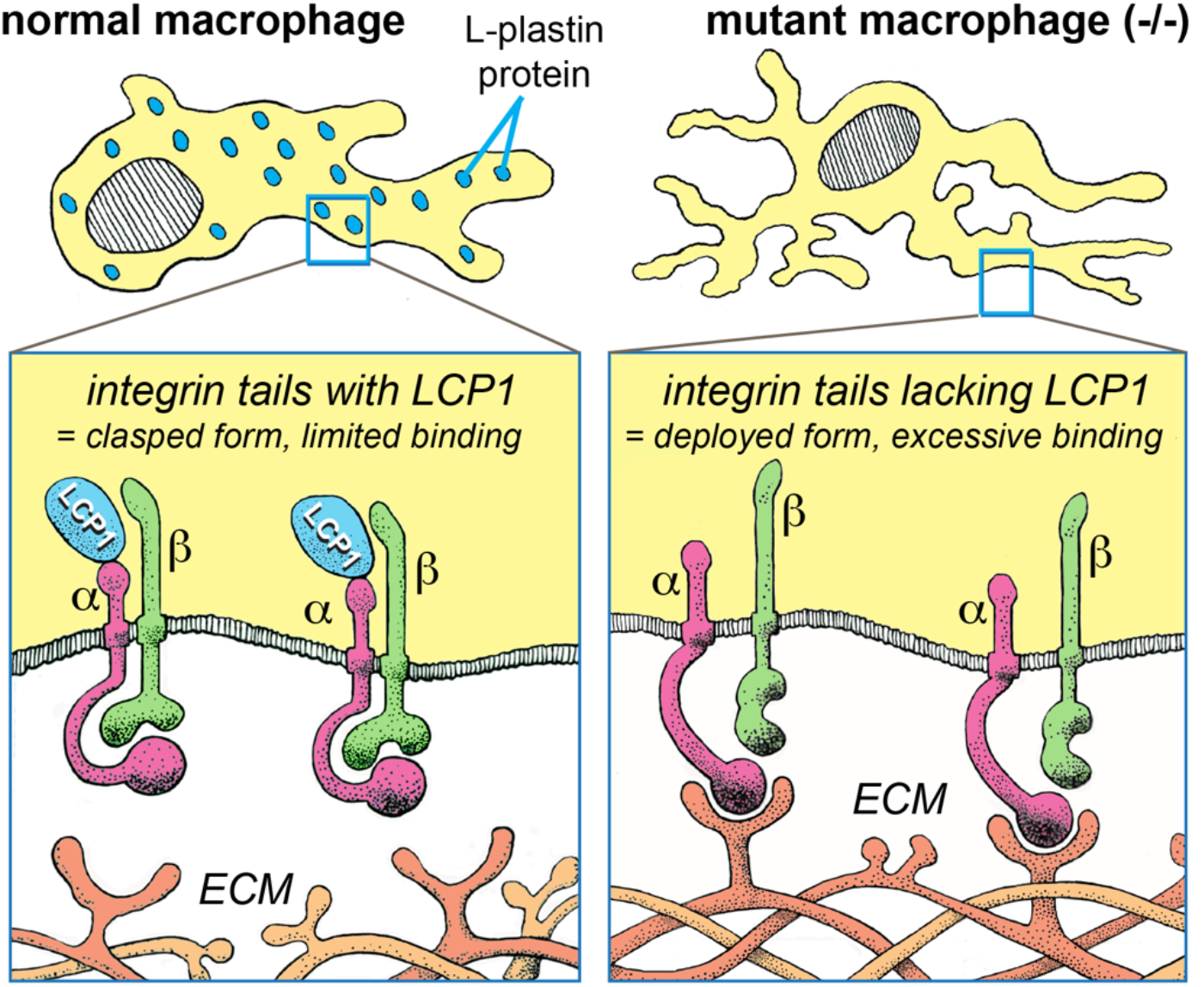
Potential role of LCP1 in regulating leukocyte integrin adhesion. Integrins are transmembrane heterodimers consisting of alpha and beta subunits (α and β). The intracellular tails (green and purple on a yellow background) are thought to interact with LCP1 (blue ovals), regulating integrin conformation and adhesion. **LEFT:** In wild type macrophages LCP1 maintains key integrins in a ‘clasped’ or low-affinity state. This reduces adhesion to the extracellular matrix (ECM) and allows cellular ‘sprinting’ in response to injury. **RIGHT**: In mutant macrophages lacking L-plastin, these integrins adopt a ‘deployed’ or high-affinity state. These are more adhesive to the ECM, inhibiting fast movement.

In contrast to the macrophage defects described above, L-plastin knockout neutrophils (*lcp1* -/-) were essentially normal in development, distribution, and migration ability. Three key findings are noted. Our first finding was that among age-matched siblings, homozygous mutant zebrafish (*lcp1*-/-) do not have a developmental size deficit at two days post-fertilization. This result is similar to observations on the L-plastin knockout mouse (LCP -/-) wherein all genotypes of sibling pups appear morphologically normal and are equally viable (Chen et al. 2003). Our second finding is that homozygous mutant larvae (*lcp1*-/-) have neither neutrophil excess (neutrophilia) nor neutrophil deficit (neutropenia) in their caudal hematopoietic tissue as compared to their heterozygote siblings (*lcp1* +/-). This finding aligns with published reports on LCP1 knockout mice showing that sibling pups of all genotypes do not differ in the overall number of circulating polymorphonuclear neutrophils, or PMN (Chen et al. 2003). Our third finding is that there was almost no difference amongst larval genotypes in the number of neutrophils drawn to a larval fin wound, the effect of genotype being plus or minus a single wandering cell. Confidence in these findings is enhanced by the fact that all larvae were from the same parents and represent full-sibling controls.

Is L-plastin therefore dispensable to neutrophil function? In this study we looked only at abundance and distribution, not biochemical activity. Interestingly, neutrophil cell lines derived from LCP1 mutant mice show decreased activation of tyrosine kinase, which is needed for the defensive ‘respiratory burst’ reaction. Therefore L-plastin deficiency might make zebrafish neutrophils less able to combat infections, and this claim could be tested using existing *in vivo* infection models in this organism (Le Guyader et al. 2008; Clatworthy et al. 2009).

A final limitation of our study is that we analyzed only two types of leukocytes– macrophages and neutrophils– within the first few days of zebrafish larval development. However, L-plastin is constitutively expressed in both innate and adaptive immune systems throughout the zebrafish life cycle (Eckfeldt et al. 2005; Bertrand et al. 2007; Feng et al. 2010). It is known that L-plastin knockout mice have poorly activated T-cells (Wang et al. 2010) and incompletely developed germinal centers (GC), the latter being required for the long-term production of B-cells (Todd et al. 2013). However, zebrafish T- and B-cells do not fully mature until several weeks post-fertilization and the effect of L-plastin deficiency on these cell types remains to be studied. Finally, L-plastin has reported effects in two other monocyte-derived cell types, namely osteoclasts and megakaryocytes. In the former it influences the cell’s adhesive and resorptive abilities (Ma et al. 2010; Chellaiah et al. 2018), and in the latter it regulates cell migration, branching, podosome formation and platelet number (Bhatlekar et al. 2020), providing other interesting contexts for the study of L-plastin function.

### Summary and Future Directions

L-plastin mutant zebrafish are available to the research community through the Zebrafish International Resource Center (ZIRC). This biological resource complements the existing L-plastin null mouse and provides many experimental advantages including large clutch size, rapid development, tissue transparency and ease of genetic manipulation. Overall, our experiments establish that *lcp1* (-/-) zebrafish have altered macrophage behavior and morphology in the context of larval injury, warranting additional studies of this association using the strengths of this biological model.

## Materials and Methods

### Fish strains and rearing

Four different zebrafish lines were used in this study. One line was the wild type ABTU strain, also called WT. The other two were the ‘Charlie’ (CH) and ‘Foxtrot’ (FX) L-plastin knockout lines (ZDB-ALT-180319-12 and 180319-13). Both lines carry premature stop codons in *lcp1,* and all homozygous mutant animals (-/-) lack L-plastin protein as previously described (Kell et al. 2018). The transgenic line Tg(*mpeg1.1*:EGFP) labels embryonic macrophages under control of a perforin-2 promoter (Spilsbury et al. 1995; Ellett et al. 2011; Ferrero et al. 2020)

All animal work was approved by the DePaul University Institute Institutional Animal Care and Use Committee (IACUC). Adult fish were housed in a 1.4–2.4 L aquaria in a recirculating housing rack filled with dechlorinated Chicago River water (pH 7.2-7.6, 28°C) with a maximum tank density of 1 fish per 200 mL. The adult diet was a 50:50 mix of finely-ground flake food (Plankton Gold Flake) and decapsulated brine shrimp eggs (Brine Shrimp Direct) once or twice daily. Fertilized eggs were obtained by natural spawning, and healthy clutches were processed for study or propagated for line maintenance. After 5–6 days of development, small fry were transferred to a stand-alone rotifer co-culture system (Best et al. 2010) for up to one week before transfer to standard housing. Larval fish (1-4 weeks post-fertilization) were fed a mixture of live rotifers and commercial powdered diet (Golden Pearl™ 50-100 and 100-300 micron sizes) until transitioned to adult food.

### Genotyping of L-plastin allele carriers

To genotype adults we used freshly-cut fin-clips processed by Proteinase K digestion in a Tween/Triton extraction buffer as previously described (Kell et al. 2018). To genotype larvae fixed in 4% paraformaldehyde-PBS and stained with Sudan Black we used a similar digestion but first heat-treated in 300 mM NaCl for 4 hours at 65°C. Later in the study we transitioned from this two-step extraction to a one-step extraction using SDS (10 mM Tris pH 8.2, 10 mM EDTA, 200 mM NaCl, 0.5% SDS, 200 ug/mL Proteinase K) following the ‘large sample number, still quick but less dirty’ genomic DNA procedure in the Zebrafish Book (Westerfield 2000). Overall, the fresh tissue protocol yielded the most consistent genotyping results. Of the two protocols for aldehyde-fixed tissue, the SDS-based extraction yielded a greater percentage of useable results (~85-90%) than the high-salt method (60-70%). Yield from the SDS-based extraction was further improved by adding 0.5 uL GlycoBlue™ per sample to the 100% ethanol step. Regardless of extraction method, genomic DNA was analyzed for L-plastin allele composition using PCR amplification and restriction digest methods previously published (Kell et al. 2018). Any embryos with inconclusive genotypes were excluded from further analysis.

### Neutrophil staining with Sudan Black

To make the staining solution, 20 mg of Sudan Black B powder (Electron Microscopy Services #21610) was mixed with 100 mL of 70% ethanol and stirred to combine. In a laboratory fume hood 200 uL of buffered phenol-chloroform-isoamyl alcohol (25:24:1) was added, and then the mixture was covered and stirred for 2 hours. After overnight settling, the supernatant was clarified through coarse filter paper, bottled, and stored at room temperature in the dark. To avoid precipitation, the solution was replaced every 6-8 weeks.

Embryos were reared in egg water containing 1-phenyl 2-thiourea (PTU) to prevent pigment development, then fixed and stained with Sudan Black (Sheehan and Storey 1947). All fixation, washing and staining steps used gentle agitation. Fixation was for 2-3 hours at room temperature in 4% paraformaldehyde / phosphate-buffered saline (PF-PBS, pH 7.4). After three brief rinses in PBST (= 1x PBS + 0.1% Tween-20), the fixed tissue was stored in 1x PBS at 4C for up to 1-2 weeks without loss of signal. Staining was for 30-40 minutes in full strength Sudan Black Solution, followed by three 10-minute washes in 70% ethanol. Ethanol washing was complete when the embryo bodies appeared white and the supernatant clear. Immediately after these washes, the embryos were rehydrated to PBST, transferred to individual wells in a 24-well plate, and stored at 4°C prior to DNA extraction and image analysis.

### Image analysis of stained neutrophils in the larval caudal hematopoietic tissue (CHT)

All images were measured blindly, without knowledge of the individual genotype. After individual image calibration, we used the polygon tool in FIJI to capture a region starting from the most posterior point of the yolk sac extension, perpendicularly across the notochord, and then around the entire margin of the caudal finfold (**Fig, 2B**). This area selection was measured in square microns (um^2^) and was used as a proxy for body size. Using the same calibrated image, we applied the Select/ Refine Edge commands in Photoshop to manually select the pixels of all Sudan Black-stained cells. After manual refinement, all selected pixels were copied to a new image, converted to 100% black, and exported to FIJI. Finally, we used the Threshold tool in FIJI to segment the binary image and return the black pixel area (um^2^).

### Larval finfold incision assay

Embryos obtained from natural spawnings were maintained in plastic Petri dishes filled with egg water at 28.5°C. The rearing solution included the anti-fungal agent methylene blue and the melanin synthesis inhibitor phenol-thio-urea (PTU). After manual dechorionation (2-3 dpf), only undamaged embryos were selected for surgery. After transfer to anesthetic water (15 parts egg water: 1 part Tricaine stock), embryos were cut across the caudal fin fold slightly distal to the posterior end of the notochord. Only 8-10 embryos were processed per batch, and all embryos in a batch were cut within 5 minutes.

To assay neutrophil migration, cut embryos were transferred to clean egg water and held at 28.5°C for 2 hours, after which they were fixed and stained with Sudan Black as described. To assay macrophage migration, cut embryos were embedded in 1% low-melt agarose in egg water (congealing temp = 28-30°C) and mounted in glass-bottomed Petri dishes (MatTek Corporation), to which was added additional anesthetic water. Embedded embryos were then transferred to the heated, humidified stage of confocal microscope for *in vivo* imaging.

### Time lapse imaging of macrophage migration

*In vivo* time lapse images were acquired using a Leica TCS SP5 confocal microscope configured to collect emissions from the Tg(mpeg1.1:Dendra2) line, in which all macrophages are labeled with a bright green fluorophore. Key microscope settings for this experiment are presented in **Table 2**. For visual reference, a single brightfield image was taken at the beginning and end of each 2-hour series.

#### Image processing and cell tracking

4D confocal stacks were loaded into FIJI and converted to multi-image TIFF files (hyperstacks). Each hyperstack was then processed using a sequence of commands including maximum projection, brightness/contrast correction and drift correction. Key settings for these major commands are given in **Table 3**. Importantly, all images were processed and measured blindly, without knowledge of the individual genotype.

Although the *mpeg1.1*:Dendra 2 transgenic line labels all macrophages, not all macrophages move towards or reach the wound. We therefore selected macrophages to track using two predetermined rules: 1) track cells obviously recruited to the wound edge and 2) track cells whose paths did not cross those of other cells. The first rule maximizes data collection from the most wound-directed macrophages, ignoring macrophages that do not move at all or show only small-scale displacements. The second rule minimizes possible tracking errors; for discussion see (Beltman et al. 2009).

#### Cell migration metrics: d, MSD, alpha and D

For each sequential image in a stack, the position of each macrophage in (x,y,t) coordinates was captured manually using Trackmate, a FIJI plugin (Tinevez et al. 2017). Cell location was defined by visual estimate of the cell’s center of mass, followed by mouse-clicking that position. In a two-person pilot study (not shown), we determined that this visual estimate had very good intra- and inter-observer repeatability, so it was used throughout our study. We then used the center of mass coordinates (x,y) extracted from each frame to generate translationally-invariant measures of cell displacement and turning angle. We define macrophage displacement (d) as the change in coordinates between two sequential frames, namely,

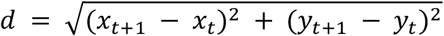

A similar calculation of displacement data was used to calculate the mean squared displacement (MSD) of each macrophage as follows:

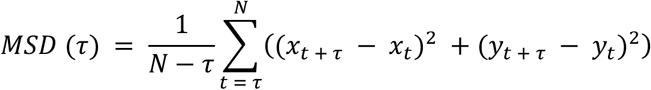

where τ (tau) represents longer intervals (1 minute, 2 minutes, 3 minutes, etc.) along a single track. Unlike displacement, which is a single number, MSD is function of time and represents the cell’s cumulative displacement as the track extends.

To calculate α and D for each track, we plot log(MSD) against log(τ). The log of a polynomial produces a linear function whose slope is D and y-intercept is α. We used the Curve Fitting toolbox in MATLAB to fit a straight line to each log-log plot and recorded the calculated slope (D), y-intercept (α) and root mean squared error (RMSE). In curve-fitting, RMSE is an estimate of the random component of the data and a RMSE statistic close to zero indicates that data fit tightly to the linear model. All RMSE were within acceptable ranges, so the linear transformation was applied.

### Null Hypothesis Statistical Testing

Individual datasets were analyzed first graphically, then statistically. Hypothesis tests were chosen as appropriate to the dataset, choosing non-parametric tests when the requirements of equal variance, normal distribution, and /or large sample size (N ≥ 20 per group) did not apply. For two-group comparisons, we used either an independent-samples Student’s t or Mann-Whitney U test to test differences in central tendency. For three-group comparisons, we used the Kruskal-Wallis test to detect differences in median value. Finally, the two-group Kolmogorov-Smirnov (KS) test evaluates global differences in distribution form.

## Acknowledgements

Financial support was provided by the National Institute of General Medical Sciences (R15-GM120664, to E. E. LeClair). Confocal analysis was made possible in part by FluoRender software funded by the National Institutes of Health (NIH R01-GM098151 and P41-GM103545).

